# Integrated molecular and functional profiling identifies E0771 as a basal-like triple-negative breast cancer model

**DOI:** 10.64898/2026.07.14.738420

**Authors:** Diana E. Baxter, Jordi Elvira-Lopez, Maria del Mar Isern, Jackeline Vanessa Huaca, Maria Teresa Blasco, Roger R. Gomis, Begoña Cánovas, Angel R. Nebreda

**Author notes:** These authors contributed equally (Diana E. Baxter, Jordi Elvira-Lopez).

## Abstract

Breast cancer is a heterogeneous disease whose clinical management relies heavily on accurate molecular subtyping. The murine E0771 mammary carcinoma cell line is widely used in preclinical studies, yet its molecular identity remains controversial, with reports variably classifying it as luminal B or triple-negative. In this study, we performed an integrated molecular and functional characterization of two independently sourced E0771 cell line stocks to resolve this discrepancy. Both stocks were genetically authenticated and exhibited concordant phenotypes. Immunohistochemical and molecular analyses demonstrated absence of oestrogen and progesterone receptors, classifying E0771 as triple-negative. Functionally, E0771 cells showed no transcriptional response to oestrogen and displayed resistance to endocrine therapy both *in vitro* and *in vivo*. Collectively, our results establish E0771 as an oestrogen-independent, basal-like triple-negative breast cancer model, supporting its appropriate use in studies of hormone-resistant breast cancer biology.

## Introduction

Breast cancer remains the most frequently diagnosed malignancy and the leading cause of cancer-related mortality among women worldwide, accounting for approximately one in four newly diagnosed cancer cases ^1^. It encompasses a highly heterogeneous group of diseases with distinct histopathological features, extensive genomic variability, and diverse clinical outcomes.

In current clinical practice, patient management is largely guided by immunohistochemistry (IHC)-based classification, which relies on the expression of the oestrogen receptor (ER) and progesterone receptor (PgR), and the over-amplification of human epidermal growth factor receptor 2 (HER2). The proliferation marker Ki67 is frequently included to distinguish luminal A from luminal B tumours, refining prognostic stratification ^2^. Based on these markers, tumours are categorized into four major subtypes: luminal A (ER⁺ and/or PgR⁺, HER2⁻), luminal B (ER⁺ and/or PgR⁺, HER2⁺ or high Ki67), HER2-enriched (ER⁻, PgR⁻, HER2⁺), and triple-negative breast cancer (TNBC; ER⁻, PgR⁻, HER2⁻). This classification informs therapeutic decisions: ER⁺ tumours are candidates for endocrine therapy, HER2⁺ tumours benefit from HER2-targeted agents such as trastuzumab, whereas triple-negative breast cancer (TNBC), lacking any of these targets, is primarily managed with cytotoxic chemotherapy ^3^.

Despite its clinical utility, IHC-based classifications only partially reflect the molecular complexity of breast cancer. To better capture its intrinsic diversity, gene expression-based classifiers were developed, most notably the Prediction Analysis of Microarray (PAM50) assay, which stratifies tumours into five intrinsic subtypes: Luminal A, Luminal B, HER2-enriched, Basal-like, and Normal-like ^4,5^. This molecular taxonomy not only recapitulates phenotypic heterogeneity but also provides independent prognostic and predictive value. Although IHC and transcriptomic subtypes broadly correlate, they do not completely overlap. For example, about 80% of TNBCs exhibit a basal-like molecular profile, underscoring the close but imperfect correspondence between these classification systems ^6–8^.

Cancer cell lines are key preclinical tools for studying tumour biology and evaluating therapeutic strategies. Extensive molecular profiling has shown that although cell lines do not fully capture the diversity of human breast tumours, they retain much of the genomic heterogeneity and recurrent copy-number alterations characteristic of the disease ^9–11^. In contrast, murine mammary tumour cell lines, which are essential for investigating tumour-microenvironment interactions and for testing therapies in immunocompetent settings, remain relatively under characterized. Several models are available that partially reflect the diversity of human breast cancer subtypes ^12^, but lack comprehensive molecular profiling, and conflicting results are often reported. Given the prognostic and therapeutic importance of molecular subtyping, a rigorous classification of murine breast cancer cell lines is crucial to ensure their biological relevance and translational value in preclinical research.

Among the available murine models, the E0771 cell line, also known as E0-771 or EO771, was originally derived from a spontaneous mammary carcinoma in a C57BL/6 mouse ^13–15^, and is widely used for studying breast cancer biology and therapeutic responses. However, the type of breast cancer that they represent remains controversial, as different studies consider E0771 cells as either luminal B ^16–18^ or triple-negative ^15,19–21^.

To address these discrepancies and determine the identity of E0771, we conducted an integrated molecular and functional characterization of E0771 cells from two different sources. By comparing their transcriptional and phenotypic profiles with an ER^+^ murine luminal control cell line, we provide evidence that E0771 cells represent an oestrogen-independent, basal-like phenotype.

## Results

### E0771 cells display features of TNBC

To address the controversy in the molecular classification of the E0771 cell line, we sourced cells from two distinct commercial providers frequently cited in the literature. Both cultures were morphologically identical, with cells exhibiting an undifferentiated spindle-shape morphology (Fig. 1A). We first performed STR analysis and confirmed a common origin for the two cell line stocks, displaying 87.5% genetic similarity to each other. When compared to the published reference standard, the stocks showed 80% and 85% identity, respectively (Fig. 1B, C). These results are consistent with cross-validation of a single cellular identity. Of note, the observed allelic mismatches were distributed across multiple loci, a characteristic pattern of genomic instability and genetic drift. The residual differences may reflect a high chromosome instability, suggesting potential variations among cell batches. Having confirmed the genetic identity of these cell lines, we proceeded with a comprehensive molecular and functional characterization to classify the E0771 subtype.

**Figure 1.**
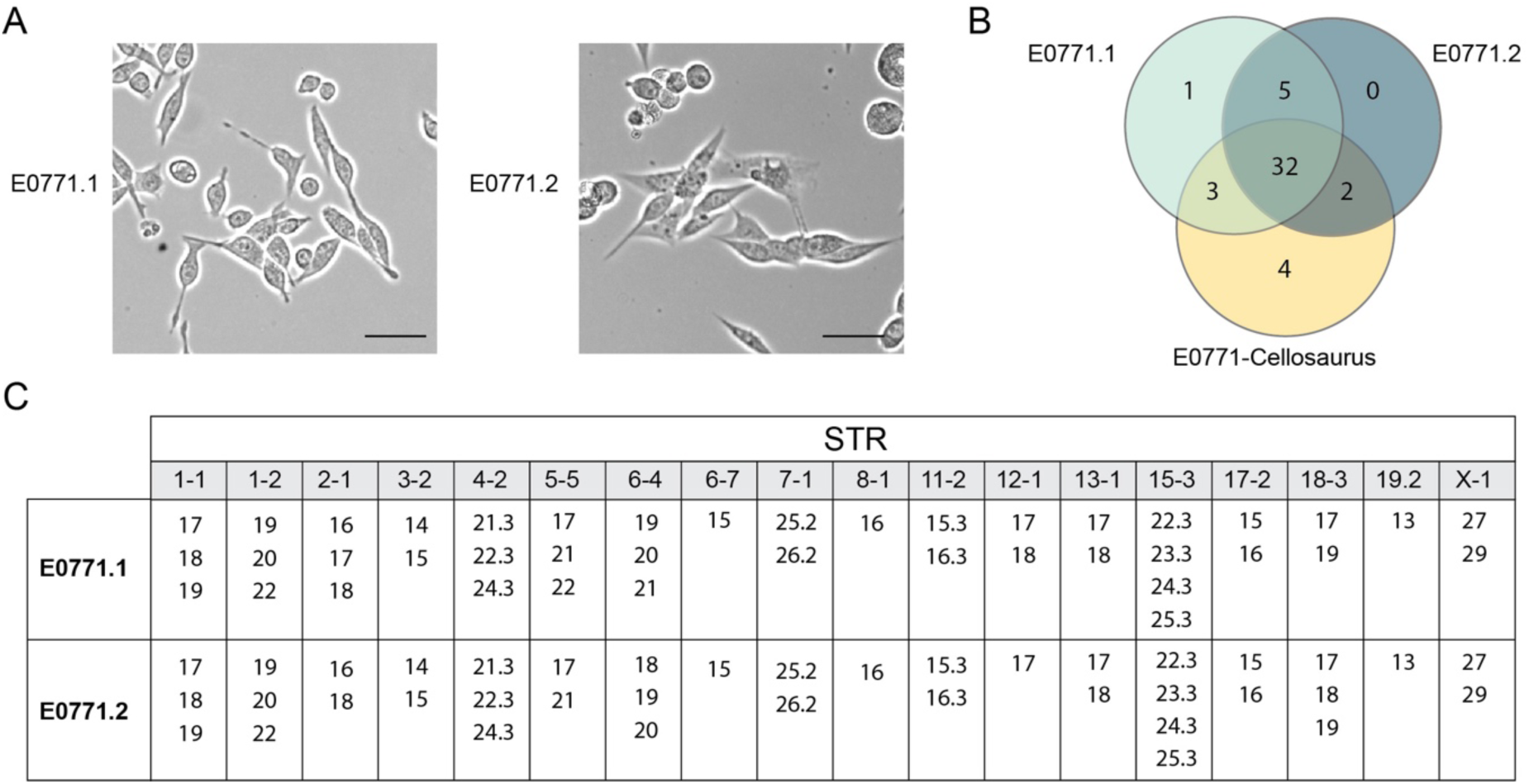
STR profiling confirms a common genetic origin for two E0771 cell line stocks. (A) Representative phase-contrast images of E0771 cells from two distinct commercial providers, E0771.1 and E0771.2. Scale bar = 50 µM. (B) Venn diagram illustrating the count and distribution of shared and unique Short Tandem Repeat (STR) alleles detected across the two commercial stocks and the published E0771-Cellosaurus reference profile. (C) Detailed table of allele calls for E0771.1 and E0771.2 across the 18 tested mouse STR loci. The numerical values indicate the repeat count for each allele identified at the corresponding locus.

In the clinical setting, breast cancer is subtyped based on the expression of three key cell surface receptors, ER, PgR and HER2. These are routinely assessed by IHC on tumour biopsy samples, where tumours are denoted as hormone receptor positive when ER and/or PgR staining is located in the nucleus. Similarly, we used IHC to assess the expression of these receptors in E0771 tumours grown in the mammary fat pad of C57BL/6 mice. E0771 tumours were negative for ER and PgR (Fig. 2A). This result classifies the E0771 cells as triple-negative, independently of the commercial source. Further analysis revealed no detectable expression of ER mRNA (Fig. 2B) or protein (Fig. 2C) in the two E0771 cell lines, which further supports the TNBC nature of this model.

**Figure 2.**
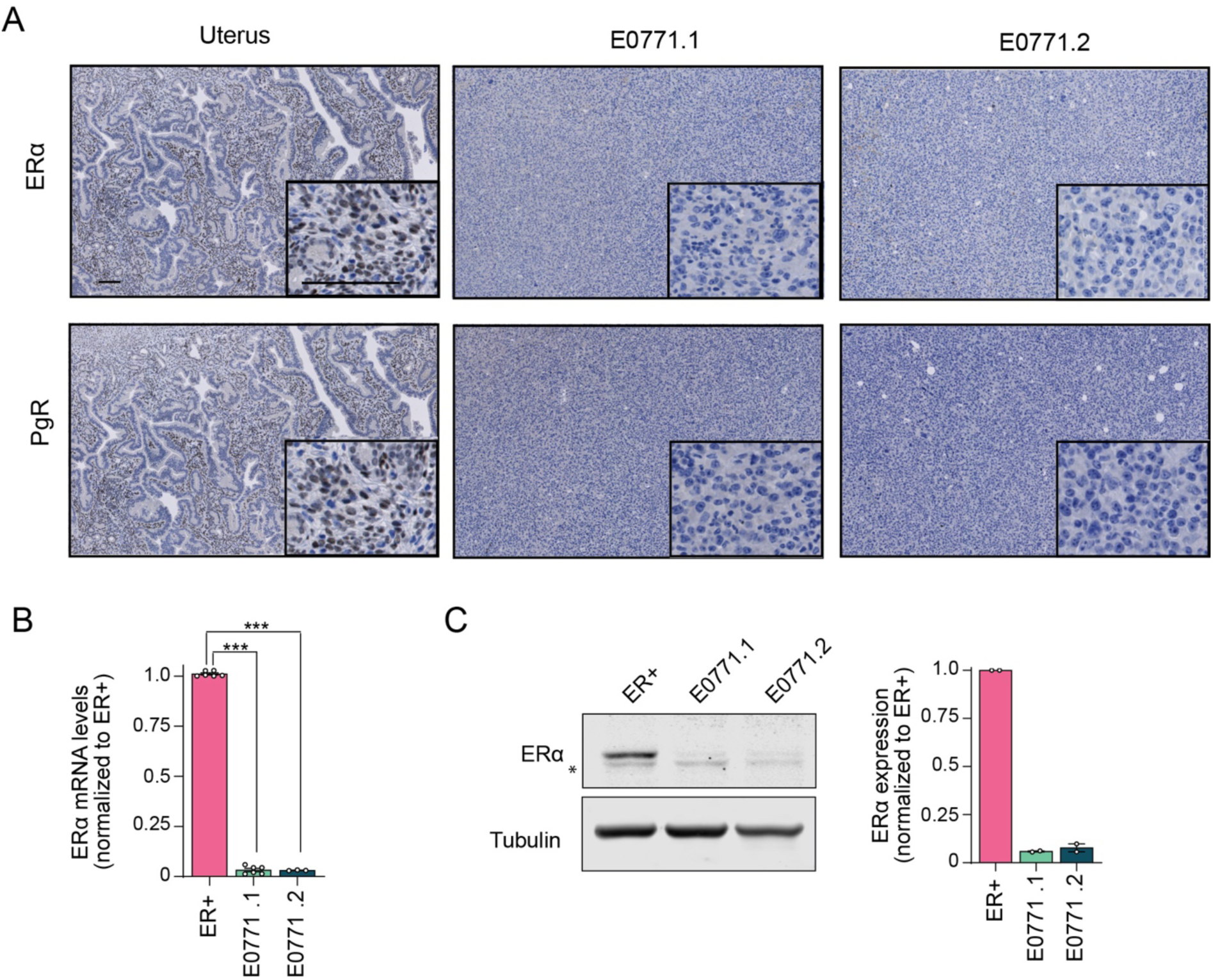
Histochemical and molecular characterization of membrane receptors classify E0771 as triple negative. (A) Representative stainings of Oestrogen Receptor α (ERα), and Progesterone Receptor (PgR) in tumours generated from the two E0771 cell stocks (E0771.1 and E0771.2). The staining shown was confirmed in four tumours. Staining of healthy mouse uterus for both markers was used as a positive control. Scale bar = 100 μm. B) Relative ERα mRNA levels in the two E0771 cell line stocks compared to a control mTB ER^+^ mouse cell line. ER^+^ n=6, E0771.1 n=5, E0771.2 n= 3. Data represent mean ± SEM. Asterisks represent statistical significance determined by Student’s t-test: *** p <0.001. C) Immunoblot showing ERα protein levels in the two E0771 cell line stocks and the mTB ER^+^ control. The asterisk indicates a non-specific band. The histogram shows the quantification of two independent experiments. Data represent mean ± SEM.

While IHC remains the clinical tool for breast cancer classification, transcriptomic profiling provides complementary information that more accurately reflects the intrinsic biological diversity of the disease. The PAM50 classifier quantifies the expression of 50 genes associated with cell proliferation, luminal differentiation, and basal identity, enabling tumours to be categorized into intrinsic molecular subtypes with distinct prognostic and therapeutic implications. To explore the molecular features of E0771 cells, we quantified the expression of all 50 PAM50 genes by qRT-PCR in the two cell lines from different sources and compared these profiles with an ER^+^ luminal mouse cell line (Fig. 3). Consistent with the STR authentication, unsupervised hierarchical clustering of the PAM50 expression matrix revealed that the two stocks of E0771 cells clustered together, confirming their genetic identity. In contrast, both E0771 cell stocks clustered separately from the mTB ER^+^ luminal cells, showing a characteristic expression pattern across the PAM50 gene set. This separation suggests that E0771 cells have a distinct transcriptional program from the luminal ER^+^ cell lineage.

**Figure 3.**
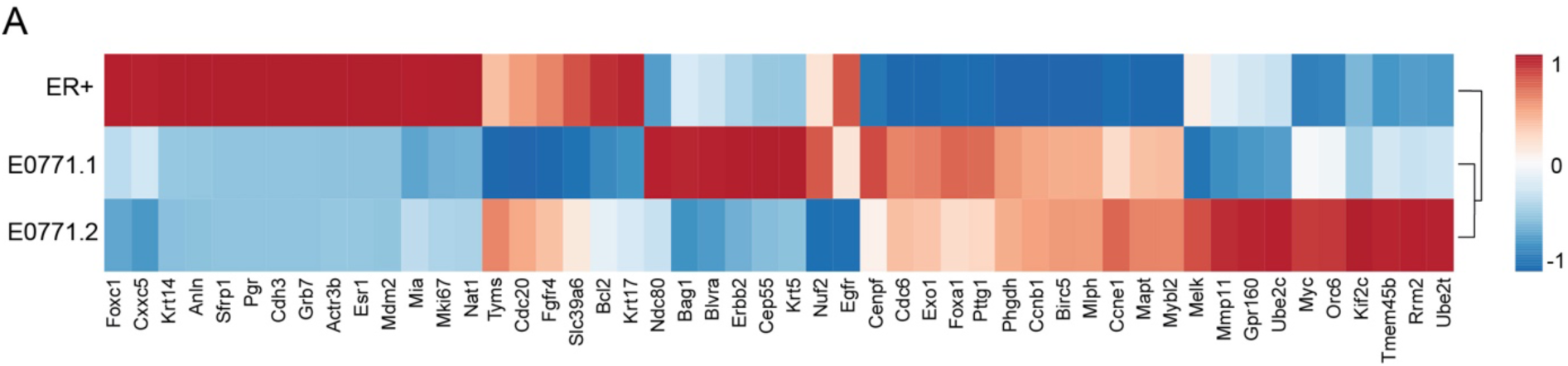
PAM50 profiling separates E0771 cells from ER^+^ luminal cells. Heatmap of the 50 PAM50 genes across the two independent E0771 cell line stocks and a mouse mTB ER⁺ luminal control. Expression values were measured by qRT-PCR in two independent samples per cell line and standardized per gene (row-wise z-scores). Unsupervised hierarchical clustering using Pearson correlation and complete linkage groups genes and samples with similar expression patterns. Colours indicate Z-score values.

### E0771 cells do not exhibit a functional transcriptional response to E2

To determine whether the IHC and transcriptional features of E0771 cells translated into functional oestrogen independence, we evaluated their transcriptional response to E2, the endogenous ligand that activates oestrogen receptor signalling. Following hormone deprivation, control mTB ER^+^ luminal cells and both E0771 cell line stocks were exposed to increasing concentrations of E2 and the expression of the canonical oestrogen-responsive genes Greb1 and Pgr was assessed after 6 h (Fig. 4A). As expected, E2 treatment robustly upregulated both Greb1 and Pgr transcripts in the ER^+^ cell line. In contrast, Greb1 was not induced by E2, and Pgr expression was undetectable in both E0771 cell stocks under all conditions tested (Fig. 4B). These findings demonstrate that E0771 cells lack functional ER signalling and are transcriptionally unresponsive to oestrogen stimulation, consistent with a triple-negative phenotype.

**Figure 4.**
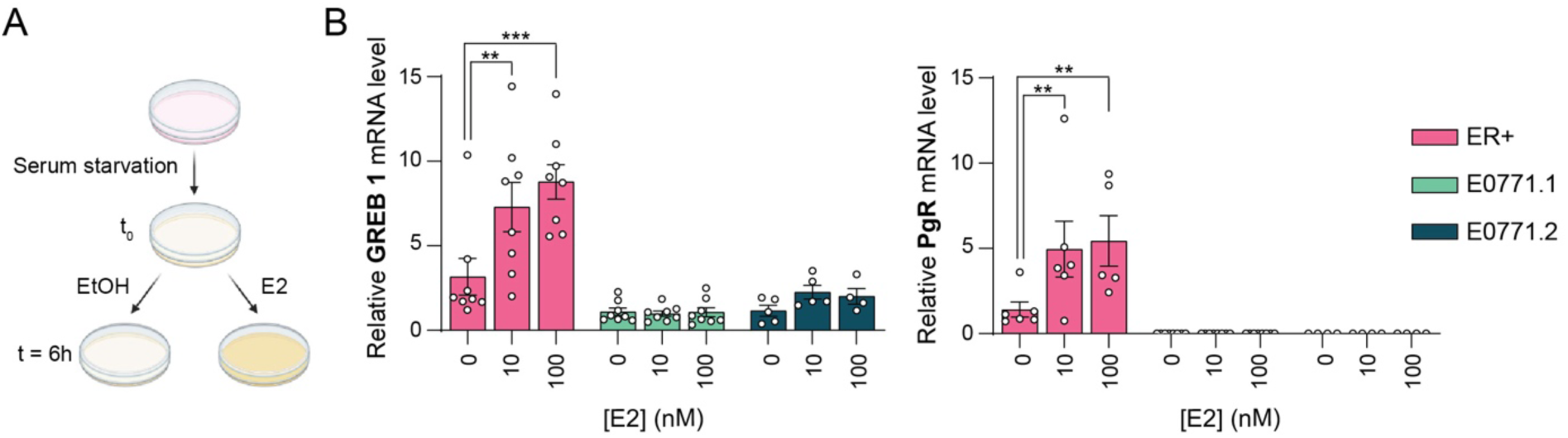
E0771 cells exhibit a lack of functional transcriptional response to oestrogen stimulation. (A) Experimental workflow for oestrogen stimulation. Mouse mTB ER^+^ luminal cells and the two E0771 cell line stocks were subjected to serum starvation (t0) followed by treatment with either vehicle (EtOH) or 17β-oestradiol (E2, 10 or 100 nM). Samples were harvested after 6 h of treatment. (B) Relative mRNA expression levels of canonical estrogen-responsive genes Greb1 (left) and PgR (right) determined by qRT-PCR. Expression levels are represented as fold change relative to t_0_. Data represent mean +/-SEM. Statistical significance was determined by two-way ANOVA followed by Bonferroni’s multiple comparisons test, comparing E2-treated groups to their respective vehicle (0 nM) controls. ** p <0.01, *** p <0.001.

### E0771 cells exhibit an endocrine-resistant phenotype in vitro and in vivo

To investigate whether the lack of hormonal responsiveness in E0771 cells translated into resistance to oestrogen receptor-targeted therapies, we performed dose-response analyses using the tamoxifen derivative 4-OHT, a potent selective oestrogen receptor modulator. We found that E0771 cells, regardless of the source, exhibited reduced sensitivity to tamoxifen compared with mTB ER^+^ luminal cells, with IC_50_ values approximately 1.5-fold higher than those observed in the ER^+^ cells (Fig. 5A, B). This difference was further accentuated in long-term clonogenic assays, which provide a more stringent measurement of therapeutic efficacy. In these experiments, E0771 cells displayed a marked resistance to tamoxifen, as evidenced by IC₅₀ values approximately an order of magnitude higher than those of the ER⁺ cells (Fig. 5C, D). Collectively, these findings indicate that E0771 cells are intrinsically refractory to endocrine therapy, consistent with an oestrogen-independent growth phenotype and further distinguishing them from canonical luminal ER^+^ models.

**Figure 5.**
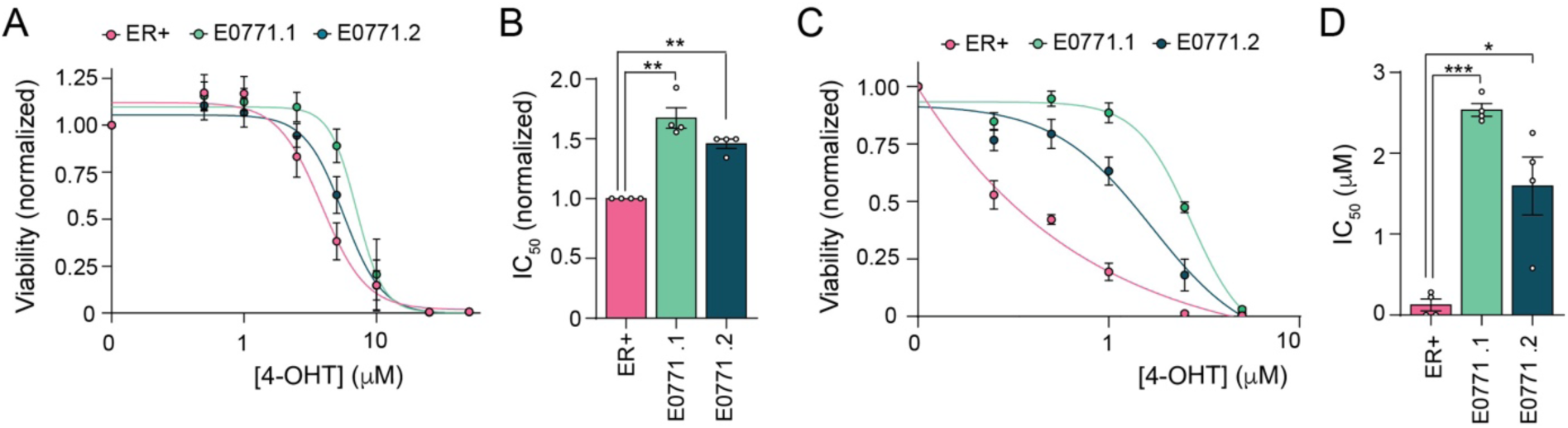
E0771 cells show reduced response to endocrine therapy *in vitro* compared to ER^+^ luminal cells. (A) Short-term dose-response curves showing normalized cell viability assessed by MTT assay following 72 h treatment with increasing concentrations of 4-OHT (n=4). (B) Histogram showing normalized IC50 values relative to mTB ER^+^ luminal cells (n=4), derived from MTT data. (C) Long-term dose-response curves showing normalized survival assessed by colony formation assay following 6-day treatment with increasing concentrations of 4-OHT. (D) Histogram showing the absolute IC_50_ values derived from the colony formation assay data. Histograms represent mean +/- SEM. Asterisks indicate statistical significance determined by Student’s t-test * p <0.05, ** p <0.01, *** p <0.001.

To determine whether the endocrine-resistant phenotype of E0771 cells was maintained *in vivo*, we compared the response to tamoxifen of tumours formed with E0771 cells or with mTB ER^+^ luminal cells. Since these models originate from distinct mouse genetic backgrounds, experiments were conducted using immunodeficient mice to avoid strain-dependent differences in tamoxifen metabolism or host responsiveness. Once tumours reached 100-150 mm³, animals were randomized to receive vehicle or 4-OHT. As expected, 4-OHT consistently suppressed the growth of the mTB ER⁺ luminal tumours (Fig. 6A), resulting in a sustained reduction in tumour size (Fig. 6B) and a markedly slower tumour growth rate, reflected by a 2-fold increase in tumour doubling time relative to vehicle-treated controls (Fig. 6C). In contrast, E0771 tumours exhibited only a slight reduction in growth at the beginning of treatment, but then the growth kinetics became indistinguishable from the vehicle-treated tumours (Fig. 6D). Although endpoint tumour volumes were modestly reduced (about 12% smaller than controls) (Fig. 6E), the overall doubling time across the treatment period remained unchanged (Fig. 6F), indicating that the early slowdown did not translate into a durable growth delay. Collectively, these data indicate that E0771 tumours retain their endocrine-resistant phenotype *in vivo*. Taken together, these findings demonstrate that E0771 cells not only diverge histologically and transcriptionally from the luminal ER⁺ cell lineage but also display a functionally oestrogen-independent and endocrine-resistant phenotype, consistent with biological features typically associated with TNBC.

**Figure 6.**
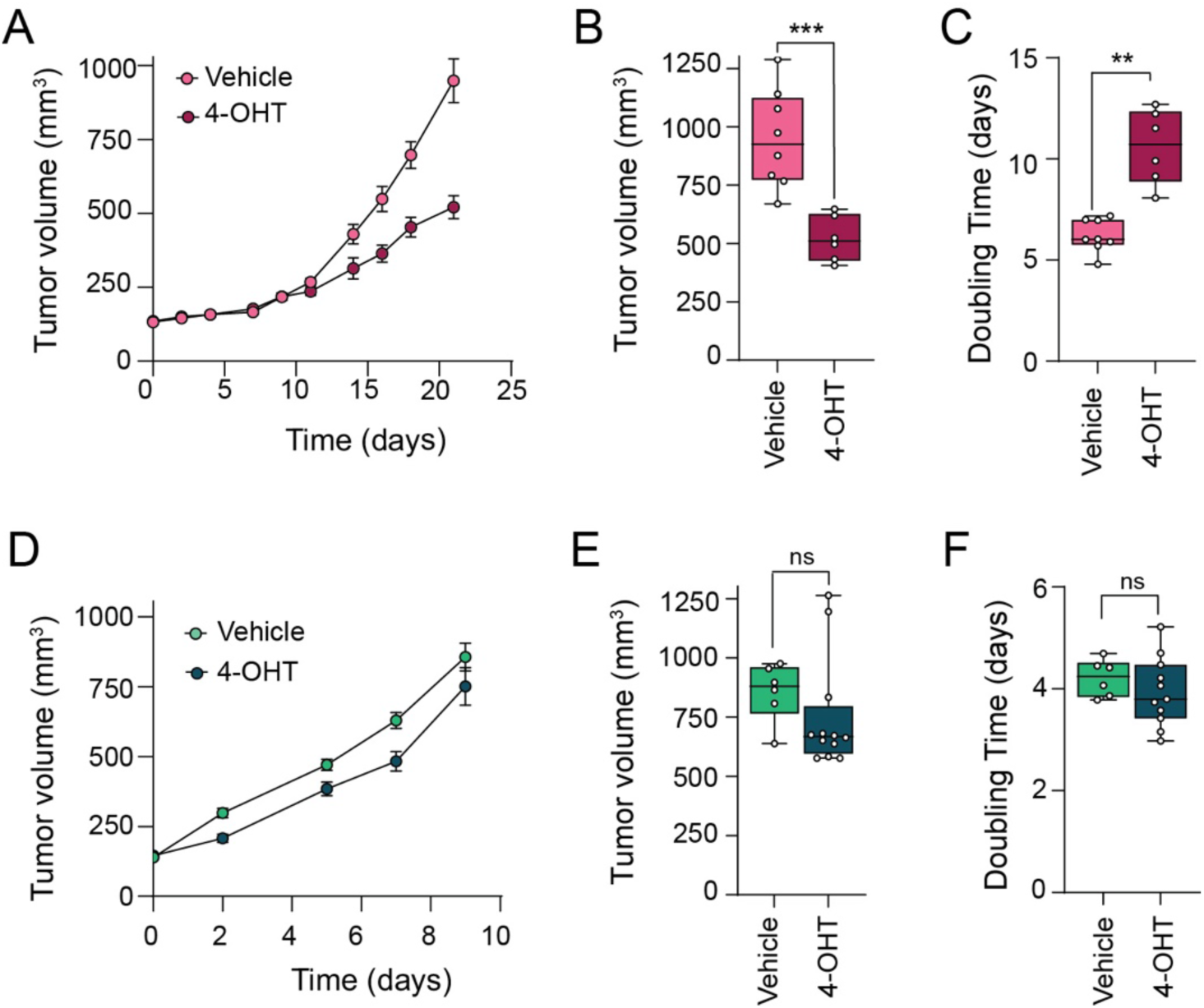
E0771 cells show reduced response to endocrine therapy *in vivo* compared to ER^+^ luminal cells. (A) mTB ER^+^ luminal cells were implanted in nude mice and when tumours were 100-150 mm^3^, mice were treated with either vehicle or 4-OHT for 16 days. Vehicle n= 8 tumours, 4-OHT n=6 tumours. (B) Final volumes of the tumours in (A). (C) Doubling growth time of the tumours in (A). (D) E0771.1 cells were implanted in nude mice and when tumours were 100-150 mm^3^, mice were treated with either vehicle or 4-OHT for 10 days. Vehicle n= 6 tumours, 4-OHT n=12 tumours. (E) Final volumes of the tumours in (D). (F) Doubling growth time of the tumours in (D). Growth curves represent mean +/- SEM. Boxplots show the median (centre line), first and third quartiles (box limits) and the minimum to maximum values (whiskers). Asterisks indicate statistical significance determined by Student’s t-test * p <0.05, ** p <0.01, *** p <0.001.

## Discussion

The success of drug development greatly relies on the translational potential of the preclinical models used. E0771 is a widely used syngeneic model in breast cancer, yet the molecular identity of this cell line remains controversial in the literature, oscillating between Luminal B and TNBC subtypes ^15–22^. Here we provide a comprehensive histological, molecular and functional characterization of the E0771 cell line using cells obtained from two main providers, benchmarking against murine mTB ER^+^ luminal cells as a control. While the lack of ER expression at the histochemical or mRNA level has been described in some publications ^15,23,24^, our results go beyond the description of biomarker expression and demonstrate the absence of functional ER signalling, which translates into a lack of response to endocrine therapy, characteristic of TNBC tumours. These findings reframe E0771 not merely as an “ER-negative” line, but as a biologically faithful model of oestrogen-independent basal-like breast cancer, validating its utility for investigating therapeutic strategies in the C57BL/6 background.

Although breast cancer subtyping and clinical ER status are defined exclusively by ERα expression, ERβ has been reported to be expressed in approximately 20% of TNBC ^25^. ERβ expression has been associated with improved survival in TNBC ^10^ and proposed to mediate limited responses to oestrogen receptor modulators through ERα-independent mechanisms ^26–28^. Consistently, E0771 have been shown to express relatively high ERβ expression ^16^. However, ERβ expression is heterogeneous and restricted to a subset of TNBC cases, and is not incorporated into current pathological guidelines ^29^ or transcriptomic classification schemes. Therefore, ERβ expression alone is likely insufficient to confer luminal identity or to classify breast cancer models as Luminal B, which requires ERα-driven transcriptional programs. These observations support the prioritization of functional interrogation of ERα pathway activity when contextualizing preclinical models of breast cancer.

Accordingly, we demonstrate that E0771 cells are refractory to oestrogen stimulation, failing to induce the canonical ER-targets Greb1 and PgR, and exhibiting resistance to tamoxifen both *in vitro* and *in vivo*. This suggests that the resistance observed in E0771 cells is a specific, cell-autonomous feature of the tumour, confirming that the model recapitulates the therapeutic resistance characteristic of human TNBC. Of note, by benchmarking these responses against a murine ER^+^ tumour cell line, rather than a human cell line, we controlled for species-specific variations in drug metabolism.

While our functional and histological data provide strong evidence for an oestrogen-independent phenotype, interpretation of molecular subtype assignments is limited by the constraints of applying PAM50 outside its original context. Despite breast cancer cell lines capturing portions of the transcriptomic landscape of patient tumours, they incompletely recapitulate lineage-specific transcriptional programs ^12^, with concordance rates of approximately 60% in human cell line panels ^10^. These limitations are likely amplified in murine models, where prior studies have classified ER-negative and TP53-mutant cell lines as luminal despite their basal-like biological features ^22^. Nevertheless, in spite of their transcriptomic variability, the two E0771 stocks used in this study consistently segregated from ERα-positive luminal controls in unsupervised clustering. This separation was driven in part by a coordinated repression of ER-associated genes, including Esr1, Pgr, Mdm2, and Nat1, supporting the absence of canonical ERα-driven luminal programs, indicating divergence from luminal identity.

It should be noted that E0771 cells display features of chromosomal instability, including elevated rates of chromosome missegregation and micronucleus formation ^30,31^, consistent with increased genomic plasticity and tumour evolution ^32^. Accordingly, STR profiling revealed genetic variability between independently sourced E0771 stocks, which goes in line with the differences we found in the transcriptional profiles and drug responses. While such genomic drift may contribute to inter-laboratory variability in growth kinetics or therapeutic sensitivity, it is unlikely to drive the loss of an established ER-driven transcriptional program or the de novo acquisition of one in a basal-like lineage.

In conclusion, our multimodal characterization classifies E0771 as a basal-like cell line. By confirming the lack of ERα and PgR expression and, more importantly, the absence of a functional oestrogen signalling axis, we validate E0771 as a robust platform for modelling TNBC.

## Methods

### Cell culture

The E0771 cell line was obtained from CH3 Biosytems (E0771.1) and the American Type Culture Collection (E0771.2). The control ER^+^ luminal cell line mTB (21154) has previously been published ^33^. E0771 and ER^+^ cells were maintained in standard conditions (37°C, 5% CO2) in Dulbecco’s Modified Eagle Medium (DMEM) (Sigma, D5796) supplemented with 10% FBS, 1% L-glutamine (Labclinics x0550-100) and 1% penicillin/streptomycin (Labclinics, L0022-100). Cells were passaged every other day and routinely tested for mycoplasma using MycoAlert Detection Kit (Lonza).

### Short Tandem Repeat (STR) analysis

STR profiling was performed by Secugen S.L. (Madrid) using a standardized mouse STR panel. Genomic DNA was isolated from harvested cells and subjected to Polymerase Chain Reaction (PCR) amplification targeting 18 specific microsatellite loci: 1−1, 1−2, 2−1, 3−2, 4−2, 5−5, 6−4, 6−7, 7−1, 8−1, 11−2, 12−1, 13−1, 15−3, 17−2, 18−3, 19−2, and X−1. PCR amplification was performed using fluorescently labelled primers to tag the microsatellite fragments. Fragment separation was achieved via capillary electrophoresis using an ABI3730 Genetic Analyzer (Applied Biosystems). The resulting fragment sizes were analysed, and genotypes were assigned using the GeneMapper software package. The STR profiles obtained from the two separately sourced cell lines were compared to each other and to the reference profile for the E0771 line available in the public Cellosaurus database using Tanabés algorithm: Similarity = 2x number shared alleles/ (number alleles cell line1 + number alleles cell line2).

### Oestrogen stimulation

2×10^5^ E0771 cells or 2.5×10^5^ ER^+^ luminal cells were seeded in 6-well plates. After 24 h, cells were placed in phenol red-free DMEM (Sigma, D1145) without FBS and 20 h later 10 nM or 100 nM of 17β-oestradiol (E2) (Sigma, E2758) were added to the media. Ethanol was used as vehicle control. Baseline, vehicle (ethanol), and E2 treated cells for 6 h were harvested for RNA extraction. To determine the relative fold change of Greb1 and PgR, all experimental groups (vehicle and E2-treated) were compared against a common baseline of cells collected immediately prior to treating with either ethanol or E2.

### Immunoblotting

Cells were resuspended in lysis buffer: 50 mM Tris HCl pH 7.5, 150 mM NaCl, 2 mM EDTA, 1% NP40, 2 mM PMSF, 2 mM mycrocystin, 2 mM sodium orthovanadate, 1 mM DTT and 1x EDTA-free complete protease inhibitor cocktail (Roche, 11873580001), mixed by vortexing and incubated 15 min on ice. Clear lysates were obtained by centrifugation at 13,200 rpm for 10 min. Cell lysates were quantified using the RC DCTM Protein Assay kit II (Bio-Rad, 5000122) with BSA as standard. Protein lysates (20 µg) were separated on 10% polyacrylamide gels and transferred onto nitrocellulose membranes (Whatman, 10401396).

Membranes were blocked in 5% non-fat milk in PBS for 1 h at RT and then incubated at 4°C overnight with antibodies against ERα (Abcam, ab241557) or Tubulin (Sigma, T9026). After washing with PBS, membranes were incubated with Alexa Fluor 680-conjugated secondary antibodies (Invitrogen 1:5000) for 1 h at RT and visualized using Odyssey Infrared Imaging System (Li-Cor, Biosciences).

### RNA extraction, reverse transcription and qPCR

Cells were washed once in Phosphate-Buffered Saline (PBS) and collected using Trizol. RNA was purified with the PureLink RNA mini-kit (Invitrogen) following manufacturer’s instructions, and quantified using a NanoDrop spectrophotometer. cDNA was obtained from 1 µg of purified RNA using SuperScript IV reverse transcriptase (Thermo Fischer Scientific).

Quantitative reverse transcription PCR (qRT-PCR) was performed in triplicates using 25 ng of cDNA mixed with 2x SYBR Select Master Mix (Applied Biosystems) and 0.2 µM of each primer in a final volume of 10 µl. Plates were run in a Quant6 Flex (Thermofisher) using the following program: 50°C for 2 min; 95°C for 10 min; 40 cycles of 95°C for 15 s and 60°C for 1 min; 95°C for 15 s; 60°C for 10 min; 95°C for 15s . Expression values were normalized to the mean of two housekeeping genes using the comparative Cycle Threshold (CT) method.

The primers used for the analysis of ER-regulated genes are indicated in Table 1, and for PAM50 profiling are shown in Supplementary Table 1.

**Table 1:**
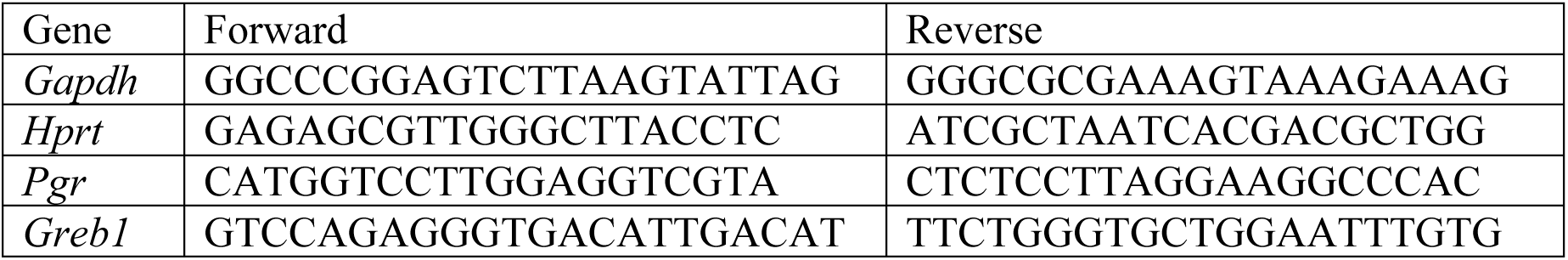
Sequences of primers used for qPCR.

### MTT assays

1,000 cells were seeded in a 96-well plate in a final volume of 100 µl and allowed to adhere for 24 h. Cells were then incubated with increasing concentrations of (Z)-4-Hydroxytamoxifen (4-OHT) (Sigma, H7904) in fresh media for 72 h. Metabolic viability was determined using the Cell proliferation Kit I (MTT) (Roche, 11465007001) following the manufacturer’s instructions. IC_50_ values were calculated by fitting the dose-response data to a four-parameter, non-linear regression model (Inhibitor vs. Response, Variable Slope) using GraphPad Prism 10 software.

### Colony formation assays

2,000 ER^+^ cells, 1,000 E0771.1 cells, and 1,500 E0771.2 cells were seeded in 6-well plates and allowed to adhere for 24 h. The following day, the medium was replaced with fresh medium containing increasing concentrations of 4-OHT. After 6 days of incubation, cells were fixed and stained with crystal violet. The total area covered by the colonies was quantified using ImageJ software and normalized to the respective untreated control for each cell line. IC_50_ values were calculated by fitting the dose-response data to a four-parameter, non-linear regression model (Inhibitor vs. Response, Variable Slope) using GraphPad Prism 10 software.

### Immunohistochemistry

Tissues were fixed overnight in 10% buffered formalin (Sigma, HT501128) and embedded in paraffin. Sections (3 μm-thick) were stained with hematoxylin and eosin (H&E). Specific antibodies were used for the following stainings: ER (Dako, M7047) and PgR (Dako, A0098). For ER staining, the signal was increased using the EnVision™ FLEX+ MOUSE LINKER (Dako, DM824). HRP-conjugated secondary antibodies (ER: Dako, DM822; PgR: ImmunoLogic, DPVR110HRP) were used and visualised using DAB (3,3-diaminobenzedine).

### Mouse experiments

To establish orthotopic tumours, 10^5^ E0771.1 cells or 10^6^ mTB ER^+^ luminal cells were resuspended in 50 μl of Matrigel (BD Biosciences) and PBS mixture (1:1 ratio) and injected into two mammary fat pads of 6-week-old female nude mice (Envigo). Additionally, mice injected with ER^+^ luminal cells were subcutaneously implanted with E2 pellets (Belma Technologies, E2-M60) to maintain plasma E2 levels necessary for tumour growth. When tumours reached 100-150 mm^3^, mice were randomized into two groups, one was left untreated and the other one received daily intraperitoneal injections of 50 mg/kg 4-OHT (Sigma, H6278) dissolved in 10% ethanol:90% corn oil. Tumour volume was calculated using the formula (length×width2)/2. Experiments were performed following the European Union, national and institutional guidelines and experimental protocols were approved by the Animal Ethics Committee of the PCB (CEEA-PCB-24-009).

### Statistical analyses

Data are represented as mean ± standard error of the mean (SEM) and statistical significance was determined using Graph Pad Prism 10 software. For mRNA expression analysis upon E2 stimulation, significance was assessed using two-way ANOVA followed by Bonferroni’s multiple comparisons test to evaluate differences between treatments within each cell line. Otherwise, statistical significance between pairs was determined using an unpaired two-tailed Welch’s t-test. p-val <0.05 (*), p-val <0.01 (**), and p-val <0.001 (***).

## List of abbreviations

4-OHT: (Z)-4-Hydroxytamoxifen
DMEM: Dulbecco’s Modified Eagle Medium
E2: 17β-oestradiol
ER: oestrogen receptor
HER2: human epidermal growth factor receptor 2
IHC: immunohistochemistry
PgR: progesterone receptor
qRT-PCR: quantitative reverse transcription PCR
TNBC: triple-negative breast cancer

## Acknowledgements

The work in our group is supported by grants from the Spanish Ministerio de Ciencia e Innovación (MICINN, PID2022-136646OB-I00, PID2022-143093OB-I00 and RED2024-153635-T), Fundación Científica de la Asociación Española Contra el Cancer (AECC, PROYE18035RODR), Fundació D’Estudis i Recerca Oncològica (FERO, BFERO2025.02) and the Catalan Agency for Management of University and Research Grants (AGAUR, 2021 SGR-909). J.E.-L. was funded by a predoctoral contract from MICINN (PRE2021-096920). We also acknowledge institutional funding from IRB Barcelona, the CERCA Programme of the Catalan Government, and the MICINN through the Centres of Excellence Severo Ochoa award.

## Author contributions

D.E.B., J.E.L. and M.dM.I. performed experiments and analyzed data, J.V.H. performed mouse experiments, M.T.B. and R.R.G. provided essential reagents and advice, B.C. and A.R.N. supervised the work, analyzed data and wrote the manuscript with contributions from all the authors.

## Data availability

All data supporting the findings of this study are available within the paper and its Supplementary Information.

## Ethical approval

Animal studies were performed following the European Union, national and institutional guidelines and experimental protocols were approved by the Animal Ethics Committee of the PCB (CEEA-PCB-24-009).

## Consent to publish

All the authors reviewed and accepted the contents of the article.

---

**Supplementary Table 1:**
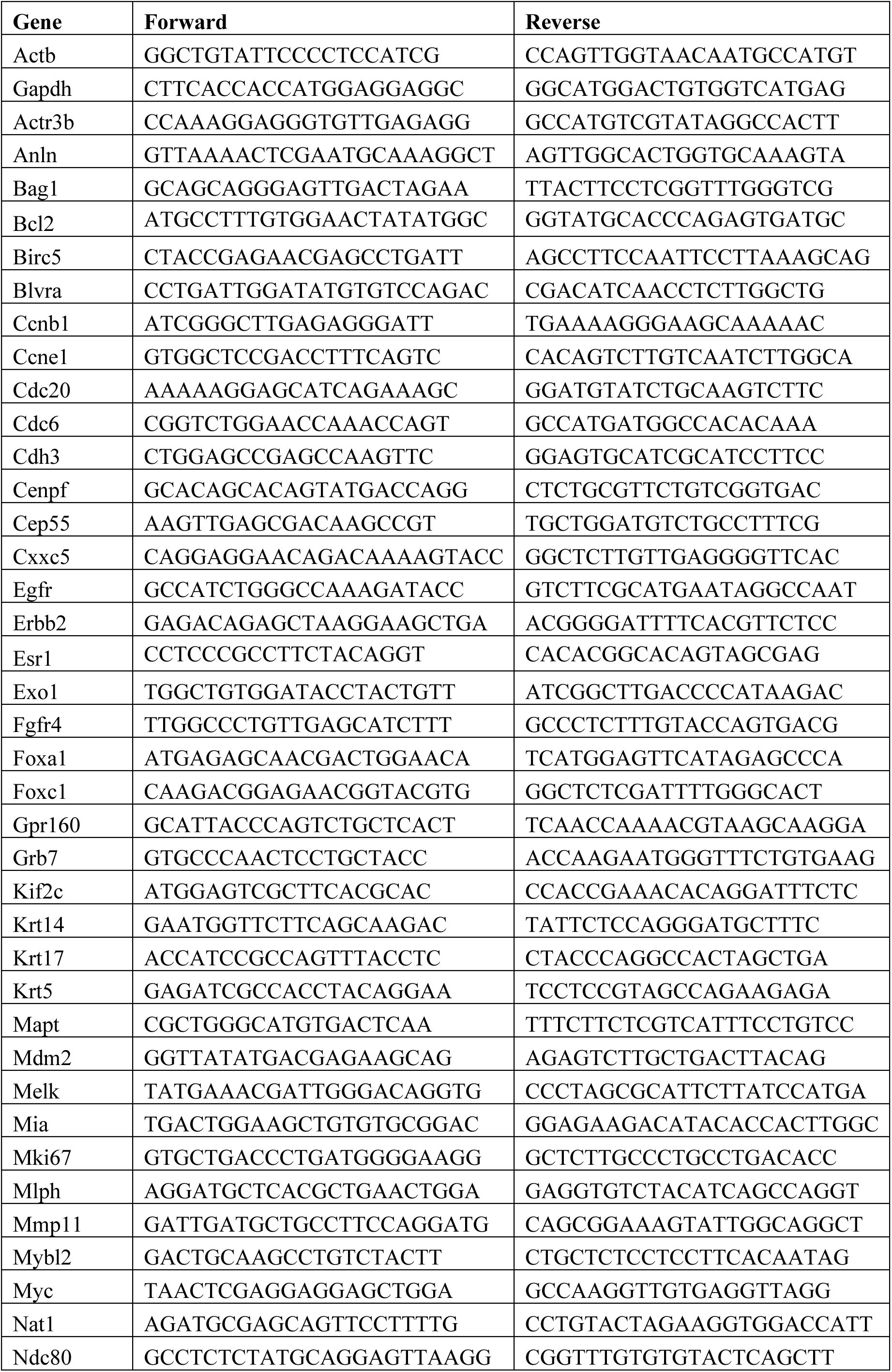
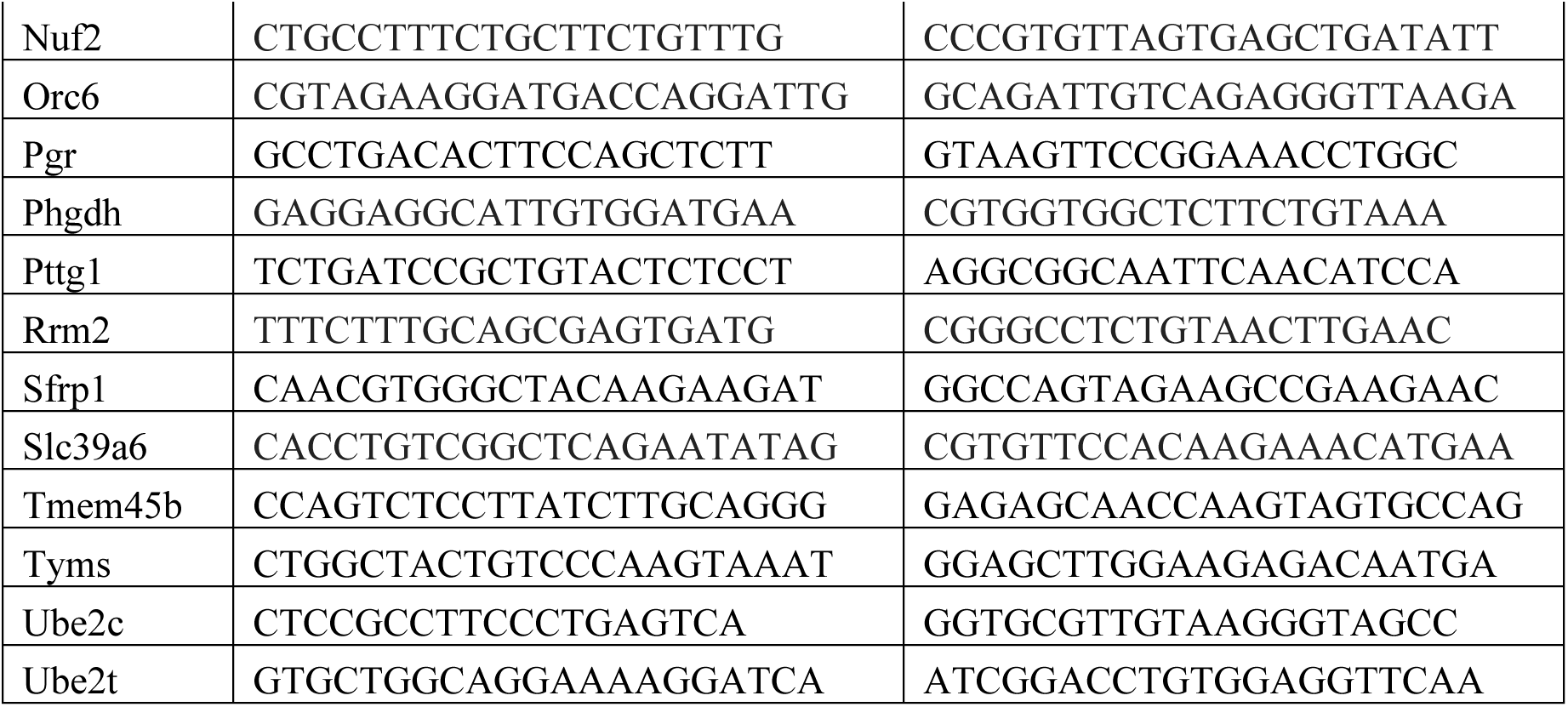
Sequences of primers used for PAM50 profiling.

